# Perturb-seq resolves physiologic programs and bidirectional regulation of non-canonical NF-κB signaling in epidermal organoids

**DOI:** 10.64898/2026.07.22.739901

**Authors:** Galen T. Squiers, Benjamin A. Nanes, Maggie M. Balas, Joshua J. Lingo, Lei Wang, Huan Zhao, Sabahat Munawar, Mpathi Nzima, Gary Hon, Jason C. Klein

## Abstract

Regulated keratinocyte differentiation is required for formation of the stratified epidermis and a functional barrier. Understanding genetic drivers of keratinocyte differentiation is crucial for understanding several skin diseases. Perturb-seq is a single-cell CRISPR screen that measures transcriptomic responses to perturbations. To date, Perturb-seq experiments have principally focused on 2-dimensional cell culture models lacking hallmarks of skin development – physiological desmosome formation and barrier function. Here, we leverage Perturb-seq in an epidermal organoid model that recapitulates physiologically relevant differentiation programs. We demonstrate that our perturbations significantly impact diverse differentiation programs and reveal bidirectional function of non-canonical NF-κB signaling in late keratinocyte differentiation.

## Background

To establish a functional barrier, basal keratinocytes progressively differentiate from a proliferative basal state to differentiated states including the spinous, granular, and cornified layers [1, 2]. This complex stratification relies on tight control of transcription factor (TF)-regulated programs, the disruption of which leads to organ malformation and disease [3-5]. Diseases of disrupted epidermal homeostasis such as ichthyoses and cutaneous squamous cell carcinoma (cSCC) pose significant global health burdens [6, 7]. To better understand the genetic underpinnings of epidermal differentiation programs, our group and others sought to identify significant drivers of keratinocyte differentiation [8]. However, most prior experiments utilized 2-dimensional (2D) models of keratinocyte differentiation which do not recapitulate crucial aspects of skin physiology and rely on multiple collected timepoints [5, 9-12]. 3-dimensional (3D) epidermal organoids (also called epidermal equivalent cultures) retain physiological properties of skin and better represent the diverse programs required for skin function [9, 13, 14]. We therefore conducted the first Perturb-seq experiments in human epidermal organoids. We profiled perturbation effects on genetic programs required for differentiation and identify a dual role for non-canonical NF-κB signaling through *NFKB2* in both the epidermal organoid and an orthogonal calcium-based differentiation model.

## Results and Discussion

We paired the well characterized air-liquid epidermal organoid with Perturb-seq and designed a library of single-guide RNAs (sgRNAs) targeting transcription factors (TFs) chosen for validated or potential roles in epidermal differentiation [15-19] (Supplementary Table 1). We plated perturbed cells to generate epidermal organoids as previously reported [5, 20, 21] and confirmed robust differentiation with hematoxylin and eosin (H&E) and canonical markers of basal and differentiated cells (Fig 1A).

**Figure 1.**
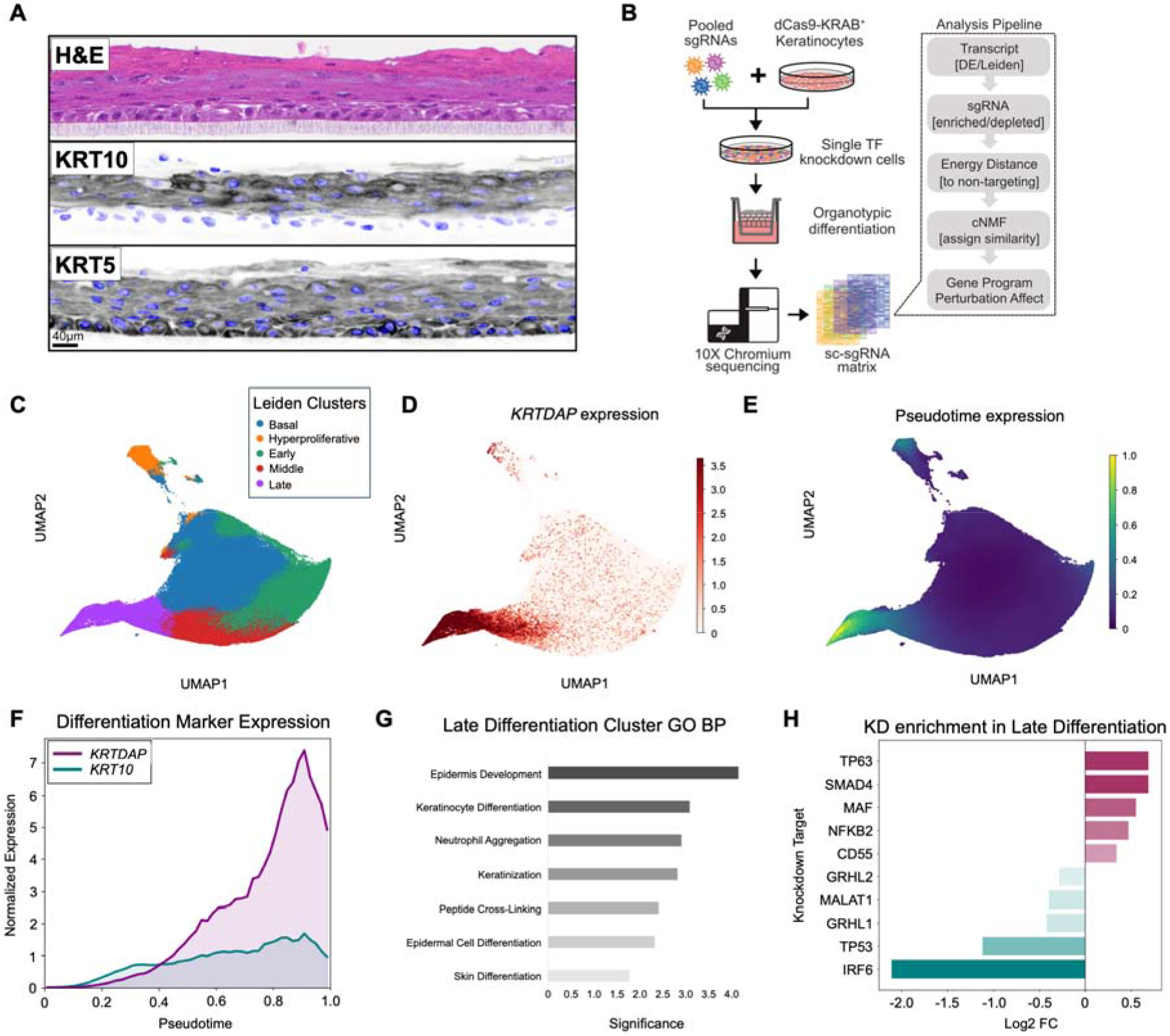
Air-liquid interface drives keratinocyte differentiation in epidermal organoids. **A)** Microscopy images of epidermal organoids stained with hematoxylin and eosin (H&E) (top), differentiated keratinocyte marker keratin 10 immunohistochemistry (IHC) (KRT10) (middle), and basal keratinocyte marker keratin 5 (KRT5) (bottom) (scale bar = 40µm). **B)** Schematic depicting Perturb-seq cell library preparation and analysis. **C)** UMAP depicting keratinocyte heterogeneity in cells isolated from epidermal organoid after 30 days of differentiation, colors correspond to Leiden clusters as shown. Each point represents one cell (n = 236,112). **D)** UMAP plot depicting normalized KRTDAP expression, color intensity corresponds to increasing expression as shown. **E)** UMAP plot depicting Pseudotime expression of cells calculated using scanpy *dpt* (root = basal). **F)** Line graph depicting *GAPDH*-normalized expression of *KRTDAP* and *KRT10* in differentiated cells over pseudotime, calculated using scanpy (root = basal). **G)** Gene ontology (GO) biological process (BP) analysis of top 100 differentially expressed genes in the late differentiation cluster (Fig. 1C, purple) calculated using *Enrichr*. **H)** Bar plot depicting Log2 fold-change (FC) of sgRNA-targets in late differentiation cluster (Fig. 2D, purple) compared to non-targeting control.

Encouraged by the robust differentiation, we dissociated perturbed cultures after differentiation and sequenced single cells (Fig. 1B).

Leiden clustering identified 5 distinct clusters corresponding to unique transcriptomic populations in the epidermal organoid [22] (Fig. 1C). Differential expression analysis identified a population expressing *KRT5, FTL* and *S100A6* corresponding to basal cells [23] (Fig. S2). Another was marked by high levels of *KRT10* and *KRTDAP* (Fig. 1C), representing differentiated keratinocytes [2, 24] (Fig. 1E, S2A). By calculating pseudotime using basal cells as the root population, we assigned average pseudotime scores per cluster as well as RNA velocity vectors for differentiation (Fig. S2B,C), revealing a progressive transition through differentiation phases. This trajectory was confirmed by increasing expression of the differentiation marker genes *KRT10* and *KRTDAP* along pseudotime in accordance with expression patterns in human skin (Fig. 1F). Gene ontology (GO) Term enrichment for the Late differentiation population highlighted keratinization and peptide cross-linking – hallmarks of spinous and granular layer development (Fig 1G). We therefore used these populations to identify preliminary perturbation effects.

We first compared the representation of TF-knockdown (KD) cells in each population and saw robust enrichment and depletion of guides in all clusters (Fig. 1H, S3A). *IRF6* is highly expressed in skin and missense mutations result in a hyperproliferative epidermis that fails to terminally differentiate [25]. We therefore expect *IRF6*-KD cells to fail to reach late differentiation stages in our model. *TP53* encodes the tumor suppressor p53, the inactivation of which is present in the majority of cSCCs [26]. *TP53-*KD cells are expected to display a hyperproliferative phenotype and fail to differentiate [27]. We confirmed that *IRF6-*targeted and *TP53*-targeted cells were depleted from the Late differentiation cluster (Fig. 1H). Interestingly, enrichment analysis showed that *NFKB2*-KD cells were among the most enriched in the Late differentiation cluster (Fig. 1H). While previous studies have focused on the contribution of canonical NF-κB signaling (principally mutations in the *NFKB1* encoded p105/p50 subunits) to epidermal differentiation [28-32], the role of non-canonical signaling via *NFKB2* in epidermal differentiation is not well described. *NFKB1* was not among the top hits in our screen (Fig. S3B), suggesting a unique role for *NFKB2* outside of the canonical signaling pathway. To validate whether *NFKB2* is indeed a negative regulator of differentiation, we sought to confirm these findings in an orthogonal system.

We treated keratinocytes with a well characterized 2D differentiation protocol that utilizes 6 days of calcium-supplemented media [1, 23, 33-35] (Fig. 2A). Wild-type 2D differentiated cells express less *NFKB2* transcript compared to undifferentiated cells, consistent with our hypothesis that NFKB2 is a negative regulator of epidermal differentiation (Fig. 2B). To validate our findings, we generated stable *NFKB2*-KD keratinocytes with shRNAs and confirmed the knockdown with RT-qPCR (Fig. S4). The *NFKB2*-KD keratinocytes have decreased expression of *CXCL14*, which we identified as a *NFKB2* responsive gene in our Perturb-seq screen (Fig. 2B, S4). Additionally, the 2D differentiated *NFKB2*-KD cells expressed more *KRT10* in both undifferentiated and differentiated cells, consistent with our hypothesis (Fig. 2D).

**Figure 2.**
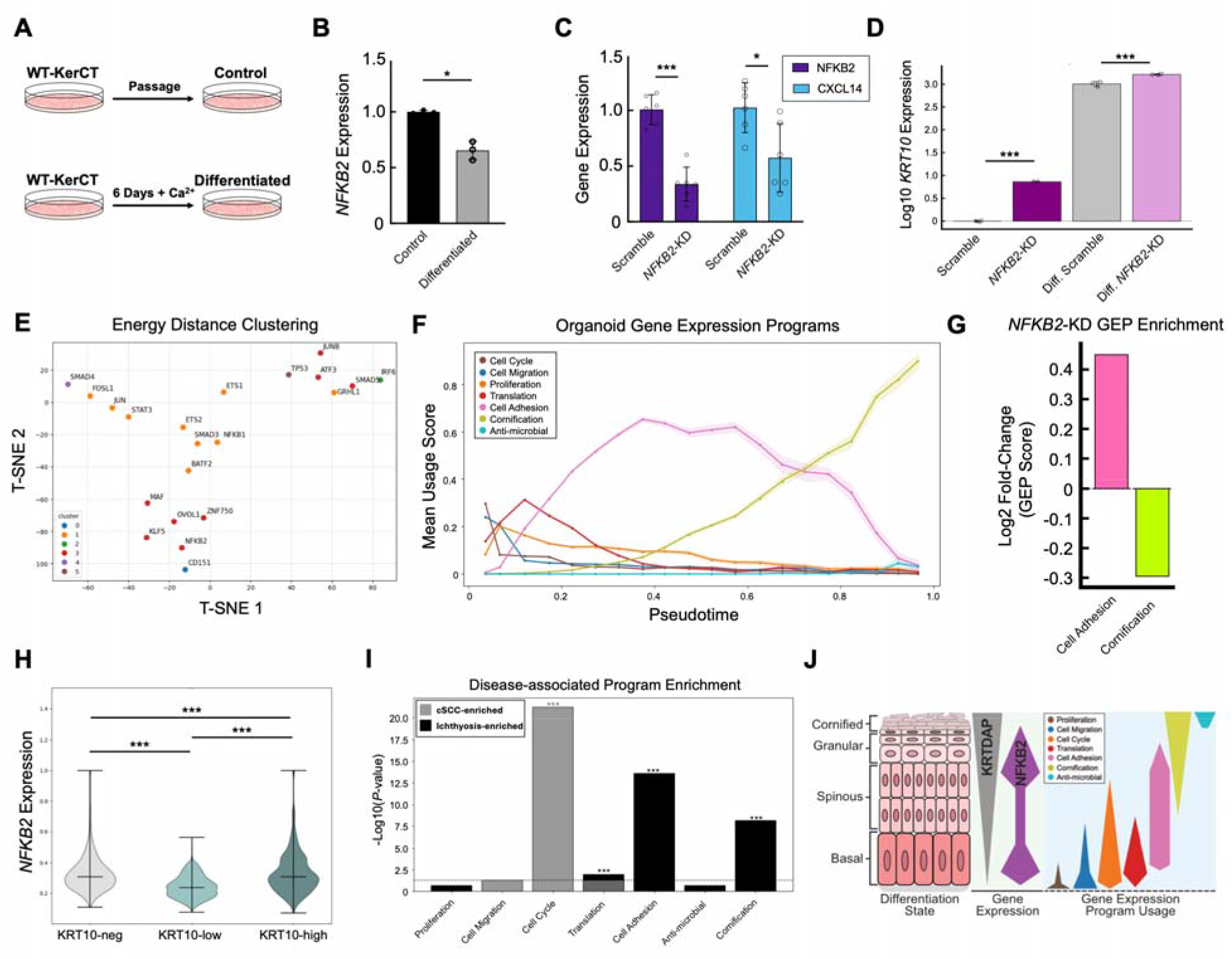
Genetic program analysis reveals perturbation and disease-specific alterations in discrete phases of differentiation. **A)** Schematic depicting calcium differentiation protocol. Wild-type KerCT (WT-KerCT) control cells were plated and passaged to prevent confluency in K-SFM growth medium. Differentiated cells were plated at confluency in 1.2 mM calcium containing K-SFM growth medium for 6 days. **B)** RT-qPCR data showing *GAPDH*-normalized *NFKB2* expression in calcium-differentiated 2D cultures (n=3). *P*-value calculated by Students t-test (* *P* < 0.05). **C)** RT-qPCR data showing expression of *GAPDH*-normalized *NFKB2* expression and *GAPDH*-normalized *CXCL14* expression in scramble control or *NFKB2*-targeted shRNA-knockdown cells from calcium-differentiated 2D cultures (n=3). *P*-value calculated by Students t-test (* *P* < 0.05, *** *P* < 0.001). **D)** RT-qPCR data showing expression of *GAPDH*-normalized *NFKB2* expression in scramble control or *NFKB2*-targeted shRNA-knockdown cells from undifferentiated and calcium-differentiated 2D cultures (n=3). *P*-value calculated by Students t-test (*** *P* < 0.001). **E)** T-SNE plot representing pairwise energy distance associations of significant perturbations from epidermal organoid Perturb-seq. 59 perturbation-target transcriptomes were filtered for significant deviation from non-targeting control (*P*-value <0.01) after which a pairwise association matrix was constructed (Energy distance = 0, no difference; Energy distance = 100, maximum difference, Fig. S5). 2D coordinates calculated via T-SNE and clusters by Affinity Propagation (*sklearn*). **F)** Gene expression program (GEP) usage in basal-to-granular keratinocyte lineage over pseudotime time represented by mean program score per pseudotime bin ± 95% CI (n = 148,499, bins = 20). Program score per cell calculated by cNMF (components = 8, iterations = 100, variable genes = 3,000). Colors represent different GEPs as indicated on graph. **G)** Log2 fold-change of *NFKB2*-KD GEP scores relative to non-targeting control cells. **H)** Violin plot showing biphasic *GAPDH*-normalized *NFKB2* expression in basal-to-granular keratinocyte lineage of epidermal organoid. KRT10-neg cells (n = 2,087). KRT10-low cells, < median expression (n = 3,019). KRT10-high cells, > median expression (n = 2,083). *P*-value calculated by Mann-Whitney two-sided t-test with Bonferroni adjustment (*** *P* < 0.001). **I)** Comparative enrichment of genes disrupted in ichthyosis and cutaneous squamous cell carcinoma, sorted by differentially expressed genes in gene expression programs. Top 30 Ichthyosis genes identified via GWAS-association (https://www.ebi.ac.uk/gwas/) [37] and cross-referenced with published disease-associated studies [38] (black). Top 50 cSCC genes identified via differential expression in NHEK vs. cSCC cell line microarray data (https://www.omicsdi.org/dataset/geo/GSE66359) [39] (grey). Top 100 genes per program calculated using cNMF. *P*-value calculated by Fisher’s Exact test with Bonferroni adjustment (*** *P* < 0.001, dotted line *P* = 0.05). **J)** Schematic overview of gene expression program utilization in different stages of basal-to-granular keratinocyte differentiation.

To explore how TF perturbations affect keratinocyte differentiation programs, we compared the transcriptomes of perturbed cells with non-targeting controls using Energy Distance scores – a measure of distance in transcriptome space [22]. This calculation identifies perturbation-associated transcriptomes that deviate from non-targeting cells (high Energy Distance). Using a p-value cutoff of 0.001 [36], we identified 21 significant perturbations (Fig. S5). We also calculated the pairwise Energy Distance between each TF-perturbed population in our library and produced six distinct clusters (Fig. 2E, S3, S5). In the T-SNE representation of these comparisons, *NFKB2* is among the highest energy distance perturbations (Fig. 2E) and we sought to explain its contributions to differentiation in the epidermal organoid.

We analyzed gene programs represented in both perturbed and non-perturbed cells using the well-characterized consensus non-negative matrix factorization (cNMF) algorithm [40-42] to 1) identify the number of gene expression program (GEP) subsets that most stably explain the transcriptional variance, and 2) assign scores for each GEP in every cell (Fig. 2F, S6A). We analyzed the top 100 variable genes for each of the 8 identified GEPs via Panther GO-enrichment and identified programs that correspond to Proliferation, Cell Migration, Cell Cycle Regulation, Translation, Cell Adhesion, Anti-microbial Function, and Cornification (Fig. S6A). We determined pseudotime vectors for each cell and plotted GEPs to correlate with differentiation state (Fig. 2F). GEP scores enhance traditional Leiden cluster analysis because each GEP represents a spectrum of expression, as opposed to a binary classification.

Because increasing Cell Adhesion GEP is implicated in the basal-spinous transition in the organoid (Fig. 2F), we sought TF-perturbations that increased Cell Adhesion GEP scores. *NFKB2*-KD cells had increased Cell Adhesion scores compared to non-targeting controls. However, upon extending our analysis to other GEPs, we saw *NFKB2*-KD cells had lower Cornification scores (Fig. 2G, S7A), counter to our hypothesis that *NFKB2* inhibits differentiation. We therefore considered a dual role for *NFKB2* in keratinocyte differentiation: one that inhibits differentiation by supporting cells in the basal state (inhibiting spinous differentiation) and another that promotes cornification. *NFKB2-*KD cells may arrest in an intermediate differentiation state; they have enhanced differentiation from basal (KRT10-neg) to spinous (KRT10-low) cells, but differentiation past this point (KRT10-high) is inhibited. Indeed, we show a significant increase in *NFKB2* in KRT10-neg (basal) and KRT10-high cells (granular) compared to KRT10-low (spinous) cells (Fig. 2G). This expression pattern also holds true when cells are segregated by *KRTDAP* expression (Fig. S7B). QuPath analysis [43] of published immunohistochemistry of NFKB2 in healthy skin [44] further supports this finding as NFKB2 is preferentially expressed in the basal (55% NFKB2+ cells) and granular (97% NFKB2+ cells) layers, with decreased expression in the spinous layer (12% NFKB2+) (Fig. S8). These data support our hypothesis that *NFKB2* inhibits early differentiation and supports late differentiation through the Cornification GEP in epidermal organoids and human skin.

Finally, we explored whether epidermal organoid GEPs are relevant to human diseases. We identified gene sets associated with skin diseases characterized by defects in cell proliferation (cutaneous squamous cell carcinoma, cSCC [6]) or barrier formation and function (ichthyosis [45]). We hypothesized that genetic drivers of these diseases would be enriched in relevant GEPs. We see genetic drivers of cSCCs are heavily enriched in Cell Cycle program-associated genes (Fig. 2I). In contrast, genetic drivers of ichthyosis are predominantly enriched in late differentiation programs such as Cell Adhesion and Cornification (Fig. 2I). In total, these data suggest that our epidermal organoid is a suitable system for massively parallel modeling of diverse disease-relevant defects. This represents a significant advancement in skin organoid methodology and analysis.

## Conclusions

In this work, we establish the first Perturb-seq screen in epidermal organoids to track the transcriptomes of perturbed cells through a gradient of differentiation states and elucidate subtle effects of TF perturbations [46]. We reveal distinct cell clusters with hallmarks of physiological differentiation and show that many of our perturbations disrupt or promote epidermal differentiation. We validate key differentiation programs and the regulator *NFKB2* in an orthogonal 2D calcium differentiation model.

Additionally, we implement a downstream analysis pipeline that provides powerful insight into perturbation mechanisms. Energy distance analysis highlights similarities between KD cells while GEP-centric analysis through cNMF identifies networks driving differentiation to explain energy distance associations. *NFKB2-*KD cells have enriched Cell Adhesion scores, consistent with KD promoting differentiation. However, *NFKB2-*KD cells have reduced Cornification scores, suggesting a second differentiation stage where *NFKB2* positively regulates late differentiation, or cornification. Further studies are needed to determine the mechanism of *NFKB2* regulation and the extent to which non-canonical NF-κB signaling contributes to the terminal stages of epidermal differentiation. Our Perturb-seq screen in epidermal organoids builds upon existing 2D models [47]. We show that expression patterns and organoid GEPs are representative of skin biology and pathology. This Perturb-seq screen opens exciting opportunities for epidermal organoid-based testing of diverse disease-associated perturbations and provides a framework for integrating perturbation-centric and gene program-centric pipelines to mechanistically study organoids beyond our model.

## Methods

### Cell culture

Human keratinocyte Ker-CT cells (ATCC CRL-4048) were developed from male human foreskin keratinocytes through expression of hTERT and CDK4 (a kind gift from Dr. Jerry Shay (UTSW Medical Center)). Keratinocytes were cultured on tissue-culture treated plastic coated with bovine type I collagen (PureCol, Advanced Biomatrix #5005). Dishes were pre-incubated in PBS containing 100 μg/mL collagen for 30-60 minutes. Sub-confluent cells were passaged in Keratinocyte serum-free medium (K-SFM) (Gibco CAT#17005042). Epidermal organoids were generated as described previously [21]. To generate the Perturb-seq cell line, human keratinocytes (KerCT: ATCC CRL-4048) were infected with dCas9-KRAB vector and single clones isolated. We generated a pooled library of sgRNAs, cloned them into pLenti-BFP vector, and packaged them in lentivirus produced with HEK293 cells (ATCC CRL-1573). HEK293s were maintained in DMEM (Gibco, CAT#11995-065) supplemented with 10% FBS and antibiotic–antimycotic reagent (Gibco #15240-062).

KerCT-dCas9-KRAB cells were infected with lentivirus at approximately 0.2 MOI and sorted on BFP expression. These cells were then plated at confluency in trans-wells. Cells were switched to Differentiation Medium (fully supplemented K-SFM mixed in equal parts with DMEM/F12 (Gibco #11320033) with 2% fetal bovine serum (FBS; Sigma F0926)) three days post-seeding and airlifted 10 days after seeding. Media was changed every three days until cultures were harvested 21 days post-seeding.

For calcium differentiation, Ker-CT cells were seeded at confluency on collagen-coated plates as described above. 1.2mM calcium-containing KSFM was added after 12 hours. Media was changed every two days and cultures harvested after six days.

All cells were maintained in a humidified incubator at 37 °C and 5% CO2.

### H&E Staining

The trans-wells containing epidermal organoids were removed from the culture dishes and incubated overnight in 4% paraformaldehyde solution at 4 °C (Electron Microscopy Sciences #15713). Samples were submitted to the UT Southwestern Histo Pathology Core Facility for embedding, sectioning, and preparation of hematoxylin and eosin (H&E) and immunohistochemistry (IHC) stained and unstained slides.

### Single-cell sequencing and Perturb-seq

Single-cell RNA sequencing was performed with 10X Chromium Single Cell Gene Expression Flex following protocol CG000632 for cell isolation. In brief, epidermal organoids were dissociated for 30 minutes at 37^°^C using Dispase, then 15 minutes at 37^°^C using Accumax. Cells were then physically triturated 10-15 times using P1000 pipette. Cell counts were performed with a Countess III FL with a DAPI stain. Cells were immediately used for library preparation per protocol “Chromium Next GEM Single Cell 3’ HT Reagent Kits v3.1 (Dual Index) with Feature Barcode technology for CRISPR Screening & Cell Multiplexing,” CG000421. One million cells per sample were used for probe hybridization. 12,000 cells were loaded per sample. Libraries were sequencing on a Novaseq X, 25B flowcell with PE 151bp reads. Additional reads were sequenced on a NextSeq2K, P3 with 28 and 90bp reads. Reads were trimmed and analyzed together with Cell Ranger Multi v7.2.0.

### Data filtering and aggregation

10X cell ranger outputs were organized, aggregated, and processed based upon the Hon Lab Perturb-Seq-Processing-Pipeline (https://github.com/Hon-lab/Perturb-Seq-Processing-Pipeline). In brief, transcript fastq files were mapped using 10X genomics Cell Ranger (reference genome: GRCh38) to produce the cell by transcriptome matrix. HTOs and sgRNAs were mapped with FBA and demultiplexed to contain only singlets with >0 sgRNAs. Finally, the multiple lanes were aggregated and combined with the Cell Ranger transcriptome matrix for downstream analysis. We isolated high confidence sgRNA-containing cells (Fig. S1) and applied a hierarchical processing pipeline to identify genetic programs driving keratinocyte differentiation [36] (Fig. 1B).

### Energy distance calculations

Energy distances were calculated using the energy distance pipeline as previously described (https://github.com/Chikara-Takeuchi/energy_dist_pipeline) [36]. P-value cutoffs for non-targeting guide comparisons were set to 0.001. T-SNE dimensions were set by pairwise-comparison.

### Pseudotime, RNA velocity, Differential Expression calculations

Pseudotime analysis was conducted on basal-to-granular lineage. Hyperproliferative cells were excluded from this analysis. Pseudotime scores calculated using *scanpy* dpt-pseudotime function [48] using the basal cell cluster as the root population. RNA velocity was calculated using *scvelo* [49]. Differential expression calculations by perturbation were conducted as previously reported using *pyspade* [50].

### cNMF and perturbation scoring

cNMF calculations were conducted based upon the tutorial as presented by the Engreitz Lab (https://github.com/EngreitzLab/cNMF_pipeline) [40, 42]. Stability was maximized at 8 components and usage programs were assigned based upon GO-term enrichment (https://geneontology.org/docs/go-enrichment-analysis/). cNMF was run for 100 iterations for the top 3000 highly variable genes.

Perturbation scores were assigned for each program per cell and the Log2 fold-change was calculated between targeting and non-targeting controls. P-value was calculated by Mann-Whitney U-test.

### Qupath analysis

Images taken from the human protein atlas [44] entry for NFKB2 in skin (HPA008422 and CAB022098) were downloaded and processed in QuPath [43]. Basal, spinous and granular layers were manually annotated. Cell and nuclear segmentation were assigned via Positive Cell Detection function with thresholds set based upon negative background cell staining.

## Table of Contents

**Supp Table 1: sgRNA dictionary and targets**

**Supp Fig 1: Quality control (QC) metrics for aggregated sequencing runs**.

**Supp Fig 2: Gene expression and pseudotime features of basal and differentiated cells in epidermal organoids**.

**Supp Fig 3: Perturbation enrichment and depletion in Leiden clusters. Supp Fig 4: *NFKB2* expression in cell subsets**.

**Supp Fig 5: Energy distance QC and metrics**.

**Supp Fig 6: cNMF expression and ranked gene expression. Supp Fig 7: Transcription factor knockdown GEP scoring. Supp Fig 8: NFKB2 expression in normal human skin**.

## Supplementary figure legends

**Supplementary Figure 1) Quality control (QC) metrics for aggregated sequencing runs. A)** Scatter plot showing UMI counts per barcoded cell, number of estimated cells per run, mean reads per cell, and median genes per cell for run LW446. Calculated via 10X Genomics Cell ranger. **B)** Scatter plot showing UMI counts per barcoded cell, number of estimated cells per run, mean reads per cell, and median genes per cell for run LW447. Calculated via 10X Genomics Cell ranger. **C)** Scatter plot showing UMI counts per barcoded cell, number of estimated cells per run, mean reads per cell, and median genes per cell for run LW448. Calculated via 10X Genomics *cell ranger*. **D)** Scatter plot showing UMI counts per barcoded cell, number of estimated cells per run, mean reads per cell, and median genes per cell for run LW449. Calculated via 10X Genomics Cell ranger.

**Supplementary Figure 2) Gene expression and pseudotime features of basal and differentiated cells in epidermal organoids. A)** Scatter plot showing top 25 ranked genes per Leiden cluster. Rank genes score calculated using *scanpy* rank genes. Global differential expression compares cluster expression to all other cells in aggregated dataset. (Clusters correspond to Leiden clusters as depicted in Fig. 1C). **B)** UMAP depicting pseudotime values in Basal-to-Granular lineage subset (hyperproliferative cluster excluded). Each point represents one cell (n = 226065). Color indicates pseudotime score as depicted in figure legend. Calculated using *scanpy* dpt-pseudotime. **C)** UMAP plot depicting pseudotime velocity and directionality in Basal-to-Granular lineage subset (hyperproliferative cluster excluded). Each point represents one cell (n = 226065). Color indicates pseudotime score as depicted in figure legend.

Calculated using *scanpy* dpt-pseudotime and *scvelo*. Arrow overlay indicates directionality of differentiation and pseudotime progression.

**Supplementary Figure 3) Perturbation enrichment and depletion in Leiden clusters. A)** Heat map depicting perturbation frequency per Leiden cluster. Log2 fold change calculated by comparing only single sgRNA-targeted knockdown (KD) cells to non-targeting control cells in each cluster. Cells grouped by target-gene as indicated on heat map. **B)** Bar graph depicting Log2 fold change of each perturbation in Late differentiation Leiden Cluster. Log2 fold change calculated by comparing only single sgRNA-targeted cells to non-targeting control cells in Late differentiation cluster. Cells grouped by target-gene as indicated on bar graph (Red = enriched KD, Blue = depleted KD). Red box indicates *NFKB2* KD. Blue box indicates *NFKB1* KD.

**Supplementary Figure 4) *NFKB2* expression in cell subsets. A)** Violin plot showing biphasic *GAPDH*-normalized *NFKB2* expression in basal-to-granular keratinocyte lineage of epidermal organoid. Error bars indicate mean ± SD. Cells segmented based upon Leiden cluster identity as indicated. *P*-value calculated by Mann-Whitney two-sided t-test (*** *P* < 0.001, * *P* < 0.05). Black bars indicate comparison. Mean, *P*-value, cell number indicated on plot. **B)** Violin plot showing biphasic *GAPDH*-normalized *NFKB2* expression in basal-to-granular keratinocyte lineage of epidermal organoid. Error bars indicate mean ± SD. Cells segmented based upon *KRTDAP* expression level, only KRTDAP expressing cells were analyzed. Cells segregated at *KRTDAP* expression median. *P*-value calculated by Mann-Whitney two-sided t-test (*** *P* < 0.001). Black bars indicate comparison. Mean, *P*-value, cell number indicated on plot. **C)** Violin plot showing biphasic *GAPDH*-normalized *NFKB2* expression in basal-to-granular keratinocyte lineage of epidermal organoid. Error bars indicate mean ± SD. Cells segmented based upon *KRT10* expression level, only KRT10 expressing cells were analyzed. Cells segregated at *KRT10* expression median. *P*-value calculated by Mann-Whitney two-sided t-test (*** *P* < 0.001). Black bars indicate comparison. Mean, *P*-value, cell number indicated on plot. **D)** Scatter plot depicting significantly altered genes in *NFKB2*-KD cells. Significance score calculated by -Log2(*P*-value), only genes with significance score > 8 included in graph. (Red = increased expression, blue = decreased expression).

Differential expression scoring calculated via *pyspade*. **E)** RT-qPCR data showing expression of *GAPDH*-normalized *NFKB2* expression and *GAPDH*-normalized *CXCL14* expression in scramble control or *NFKB2*-targeted shRNA-knockdown cells from undifferentiated control cultures (n=3). *P*-value calculated by Students t-test (** *P* < 0.01).

**Supplementary Figure 5) Energy distance QC and metrics. A)** Bar plots indicating the number of perturbations (sgRNA species) excluded from energy distance analysis. **B)** Heat map indicating number of perturbations (target-genes) at *P*-value and energy distance thresholds. Color corresponds to perturbation number as indicated on heat map. *P*-value of 0.001 was selected for further analysis. **C)** Scatter density plot depicting sgRNA *P*-values as a function of transcriptome distance (Log10 Energy Distance). Each point represents a single perturbation (target-gene). **D)** Heat-map depicting pairwise Energy Distance between perturbations (target-gene). Color represents pairwise distance as indicated on heat map.

Clustered genes are grouped and labeled based upon top go enrichment terms, or by perturbation name if only including a single perturbation.

**Supplementary Figure 6) cNMF expression and ranked gene expression. A)** UMAP plots depicting cNMF calculated gene expression program (GEP) scores per cell. Each point represents one cell. Color represents expression score as indicated on plot. Each plot represents the score of one GEP in the entire population. **B)** Dot plot depicting top 10 ranked genes expressed in high GEP scoring cells. Cells segregated based on top GEP score per cell. Rank genes calculated using *scanpy* rank genes. GEP group and gene expression indicated on plot. Dot size corresponds to population percentage expressing the indicated gene. Color indicates normalized gene expression in the population.

**Supplementary Figure 7)** Transcription **factor knockdown GEP scoring. A)** Bar plot depicting Log2 fold change of GEP scores in *NFKB2*-KD cells compared to non-targeting control cells. GEP identity indicated on graph, ordered by median pseudotime. **B)** Violin plot showing biphasic *GAPDH*-normalized *NFKB2* expression in basal-to-granular keratinocyte lineage of epidermal organoid. Error bars indicate mean ± SD. Cells segmented based upon *KRTDAP* expression level. Cells segregated at *KRTDAP* expression median. *P*-value calculated by Mann-Whitney two-sided t-test (*** *P* < 0.001). Black bars indicate comparison. Mean, *P*-value, cell number indicated on plot.

**Supplementary Figure 8) NFKB2 expression in normal human skin. A)** Schematic depicting annotation and quantification workflow for NFKB2 IHC analysis. **B)** Region of interest (ROI) for human protein atlas (HPA) sample HPA008422. **C)** QuPath cell percentage expressing NFKB2 in ROI for HPA sample CAB022098. **D)** ROI for HPA sample HPA008422. **E)** QuPath cell percentage expressing NFKB2 in ROI for HPA sample CAB022098.

## Declarations

### Ethics approval and consent to participate

Not applicable

### Consent for publication

Not applicable

## Availability of data and materials

The datasets used and/or analyzed during the current study are available from the corresponding author on reasonable request.

## Authors’ contributions

G.T.S. and J.C.K. analyzed Perturb-seq screen, prepared manuscript. G.T.S. generated calcium differentiated keratinocyte cultures and analysis. B.N. and S.M. prepared epidermal organoids, generated H&E and IHC data, and generated *NFKB2*-KD cell line. M.B. and J.J.L. prepared manuscript elements and generated qPCR data. J.C.K., L.W., and H.Z. generated Perturb-seq library and perturbed Ker-CT cell line. M.N. and G.H. guided Perturb-seq screen analysis and established pipelines for screen QC and downstream validations. All authors read and approved the final manuscript.

## Acknowledgments

We would like to thank the labs of Dr. Wang and Dr. Shellman for insightful conversations and feedback in the preparation of this manuscript, Dr. Takeuchi and Dr. Sundarrajan for their assistance and troubleshooting of bioinformatic analysis.

