## Supplemental Figures 1-8 for "Perturb-seq resolves physiologic programs and bidirectional regulation of non-canonical NF-κB signaling in epidermal organoids"

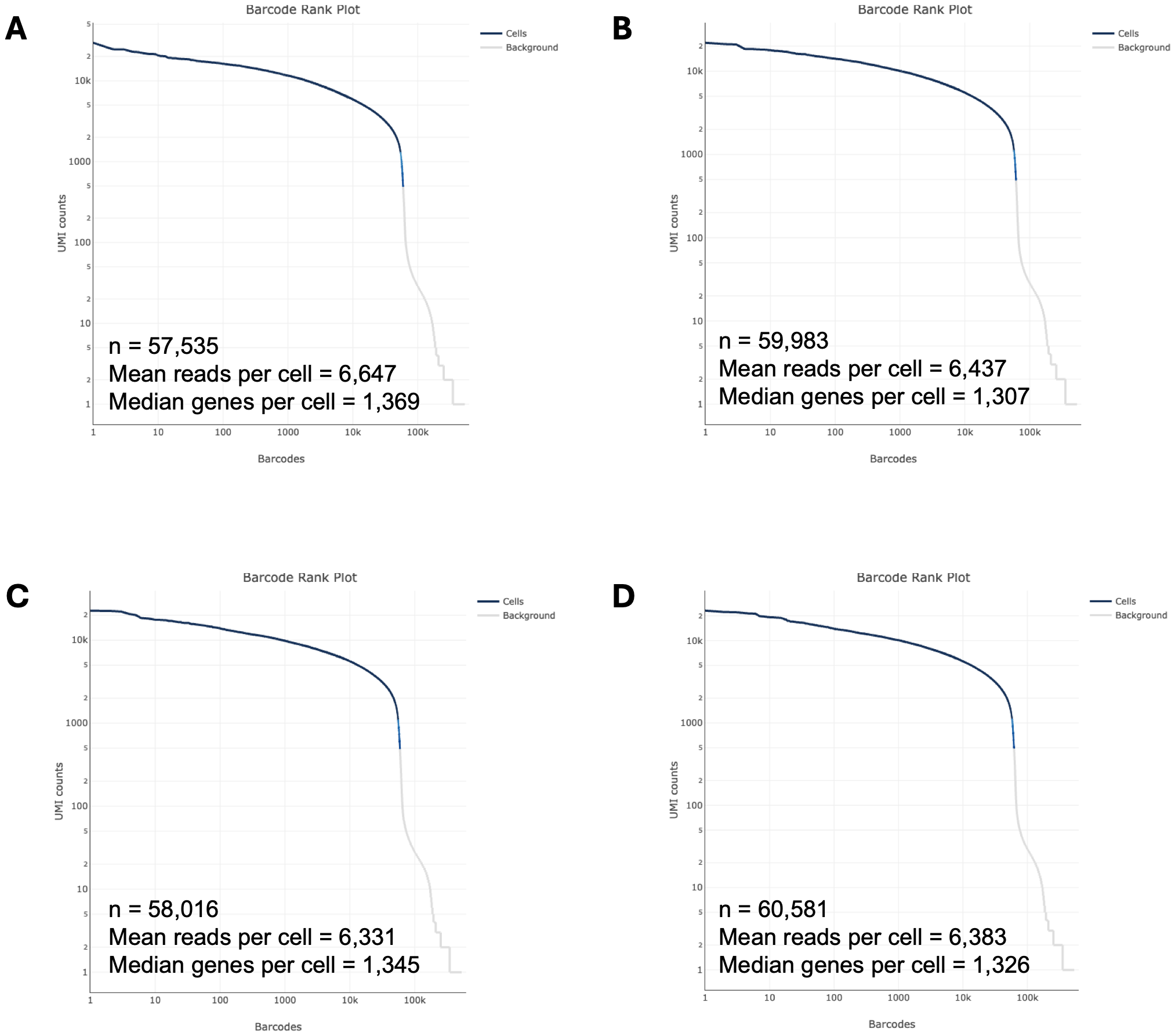

**Supplementary Figure 1) Quality control (QC) metrics for aggregated sequencing runs. A)** Scatter plot showing UMI counts per barcoded cell, number of estimated cells per run, mean reads per cell, and median genes per cell for run LW446. Calculated via 10X Genomics Cell ranger. **B)** Scatter plot showing UMI counts per barcoded cell, number of estimated cells per run, mean reads per cell, and median genes per cell for run LW447. Calculated via 10X Genomics Cell ranger. **C)** Scatter plot showing UMI counts per barcoded cell, number of estimated cells per run, mean reads per cell, and median genes per cell for run LW448. Calculated via 10X Genomics *cell ranger*. **D)** Scatter plot showing UMI counts per barcoded cell, number of estimated cells per run, mean reads per cell, and median genes per cell for run LW449. Calculated via 10X Genomics Cell ranger.

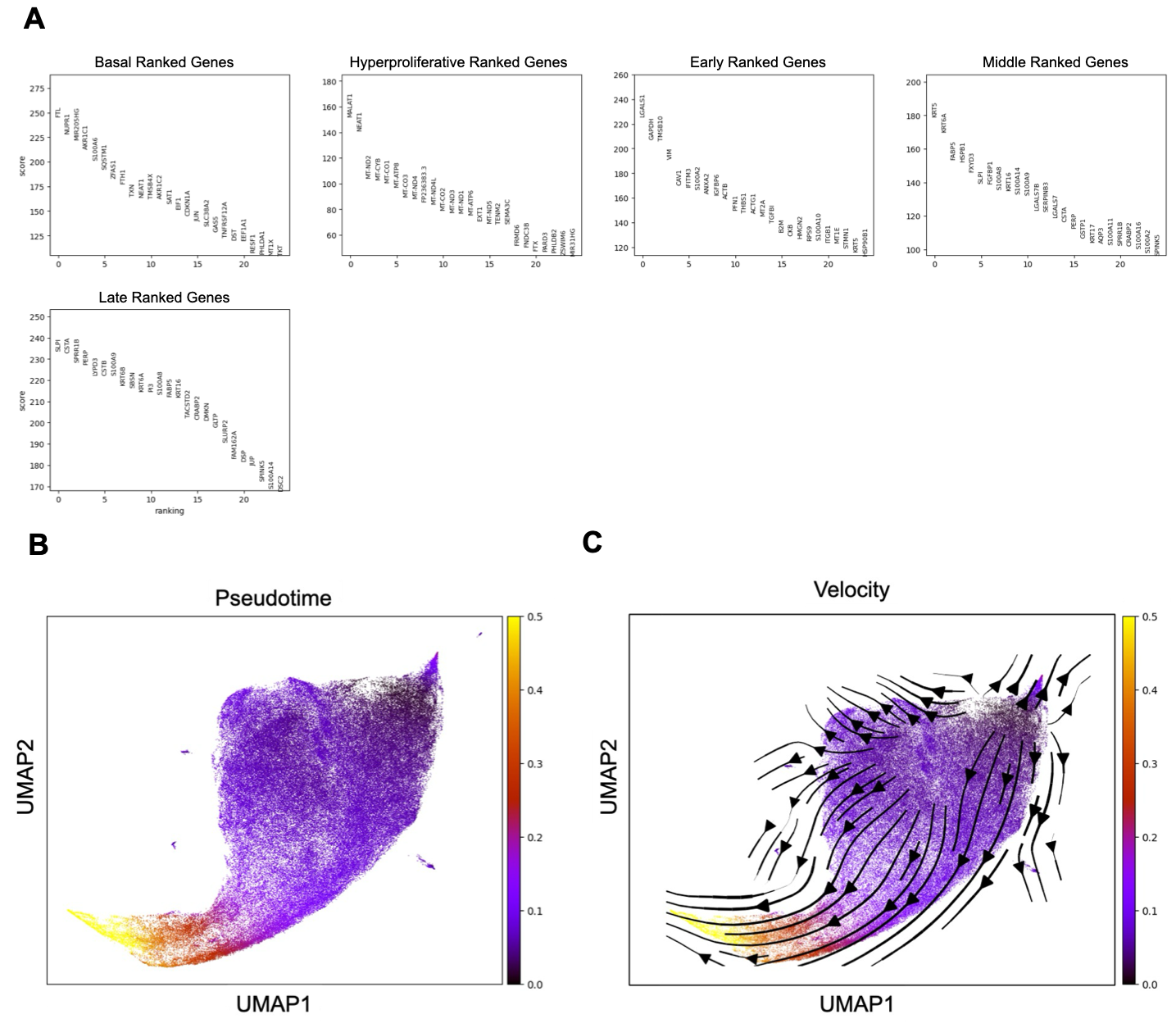

**Supplementary Figure 2) Gene expression and pseudotime features of basal and differentiated cells in epidermal organoids. A)** Scatter plot showing top 25 ranked genes per Leiden cluster. Rank genes score calculated using *scanpy* rank genes. Global differential expression compares cluster expression to all other cells in aggregated dataset. (Clusters correspond to Leiden clusters as depicted in Fig. 1C). **B)** UMAP depicting pseudotime values in Basal-to-Granular lineage subset (hyperproliferative cluster excluded). Each point represents one cell (n = 226065). Color indicates pseudotime score as depicted in figure legend. Calculated using *scanpy* dpt-pseudotime. **C)** UMAP plot depicting pseudotime velocity and directionality in Basal-to-Granular lineage subset (hyperproliferative cluster excluded). Each point represents one cell (n = 226065). Color indicates pseudotime score as depicted in figure legend. Calculated using *scanpy* dpt-pseudotime and *scvelo*. Arrow overlay indicates directionality of differentiation and pseudotime progression.

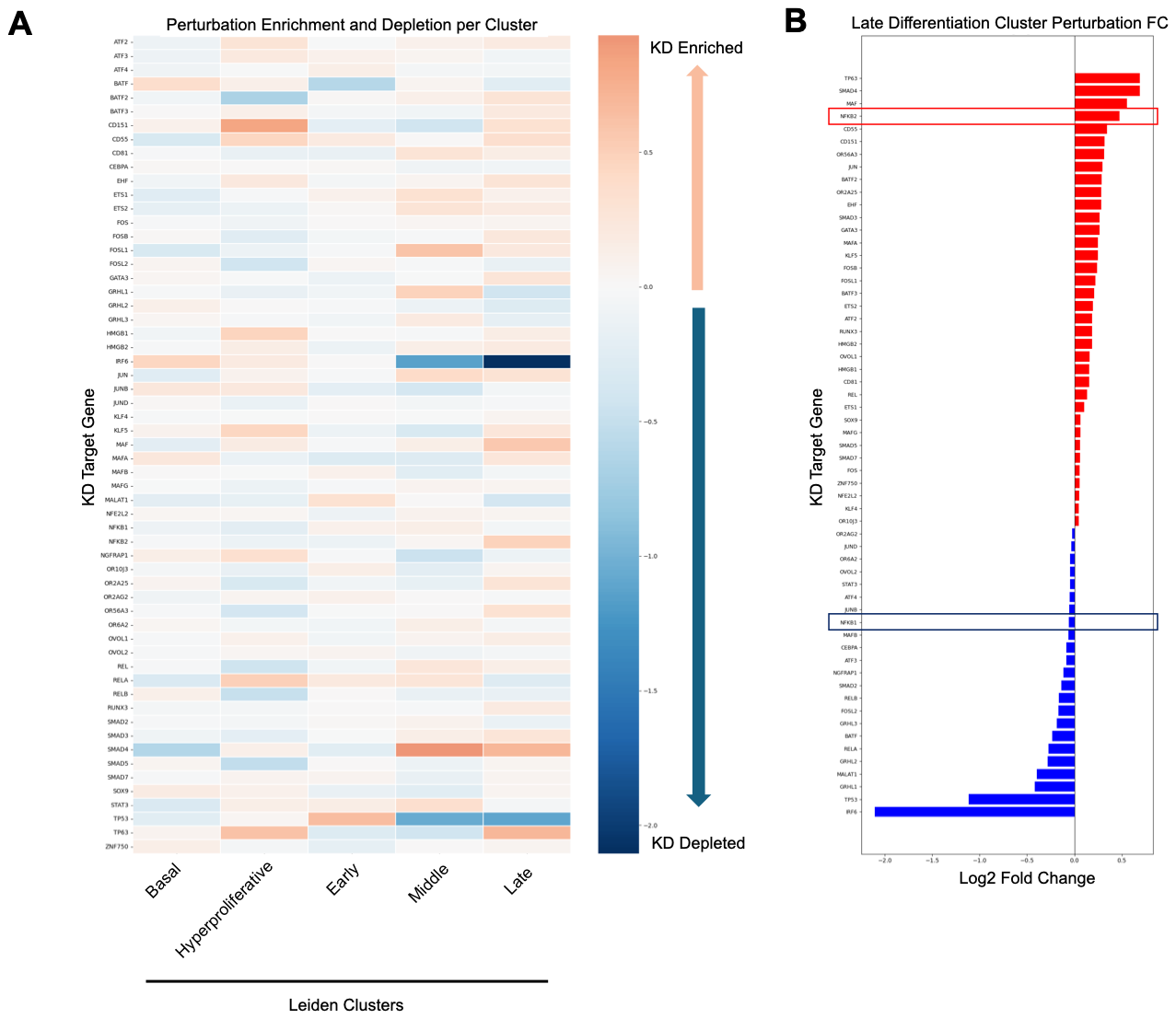

**Supplementary Figure 3) Perturbation enrichment and depletion in Leiden clusters.** **A)** Heat map depicting perturbation frequency per Leiden cluster. Log2 fold change calculated by comparing only single sgRNA-targeted knockdown (KD) cells to non-targeting control cells in each cluster. Cells grouped by target-gene as indicated on heat map. **B)** Bar graph depicting Log2 fold change of each perturbation in Late differentiation Leiden Cluster. Log2 fold change calculated by comparing only single sgRNA-targeted cells to non-targeting control cells in Late differentiation cluster. Cells grouped by target-gene as indicated on bar graph (Red = enriched KD, Blue = depleted KD). Red box indicates *NFKB2* KD. Blue box indicates *NFKB1* KD.

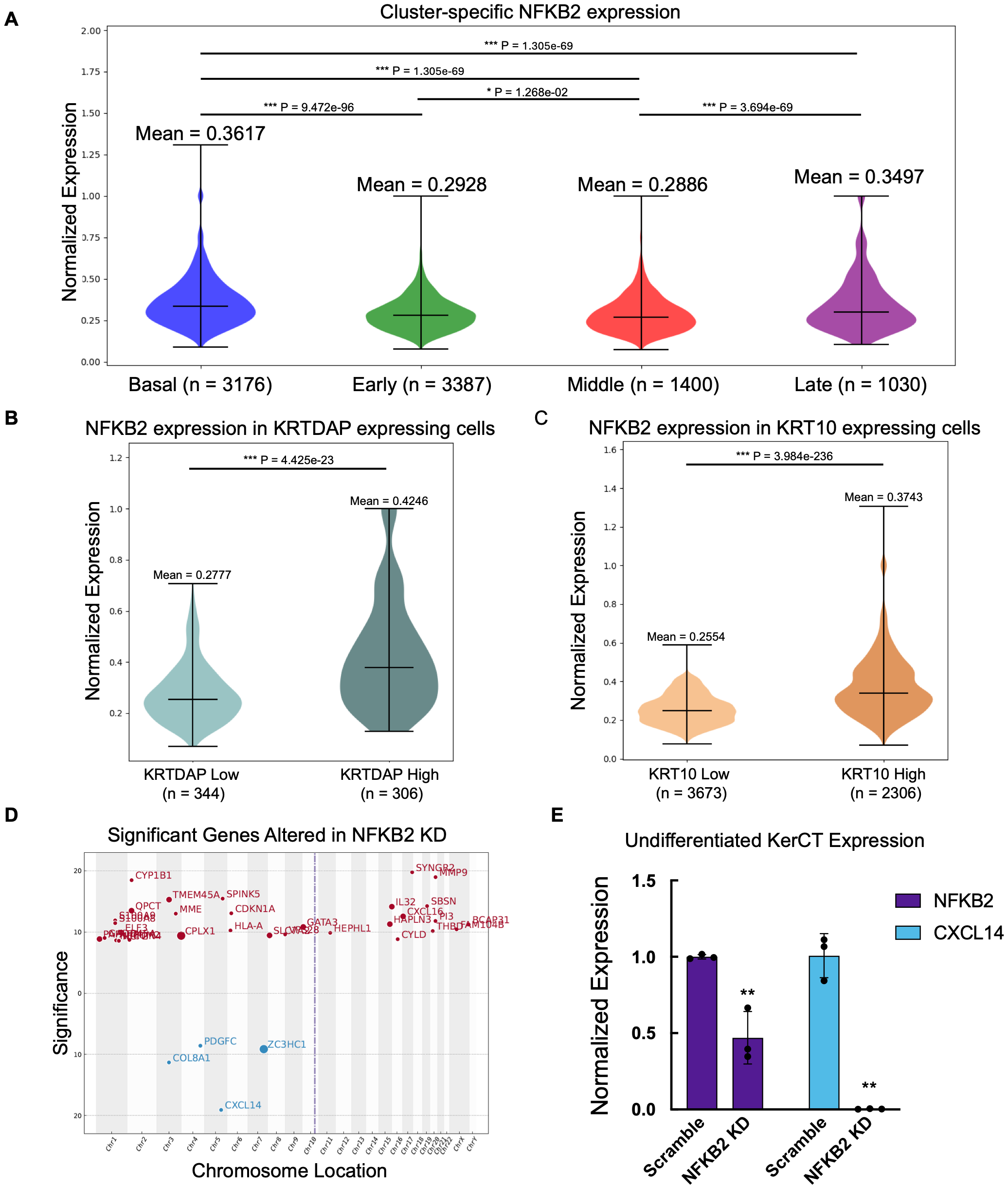

**Supplementary Figure 4) *NFKB2* expression in cell subsets. A)** Violin plot showing biphasic *GAPDH*-normalized *NFKB2* expression in basal-to-granular keratinocyte lineage of epidermal organoid. Error bars indicate mean ± SD. Cells segmented based upon Leiden cluster identity as indicated. *P*-value calculated by Mann-Whitney two-sided t-test (*** *P* < 0.001, * *P* < 0.05). Black bars indicate comparison. Mean, *P*-value, cell number indicated on plot. **B)** Violin plot showing biphasic *GAPDH*-normalized *NFKB2* expression in basal-to-granular keratinocyte lineage of epidermal organoid. Error bars indicate mean ± SD. Cells segmented based upon *KRTDAP* expression level, only KRTDAP expressing cells were analyzed. Cells segregated at *KRTDAP* expression median. *P*-value calculated by Mann-Whitney two-sided t-test (*** *P* < 0.001). Black bars indicate comparison. Mean, *P*-value, cell number indicated on plot. **C)** Violin plot showing biphasic *GAPDH*-normalized *NFKB2* expression in basal-to-granular keratinocyte lineage of epidermal organoid. Error bars indicate mean ± SD. Cells segmented based upon *KRT10* expression level, only KRT10 expressing cells were analyzed. Cells segregated at *KRT10* expression median. *P*-value calculated by Mann-Whitney two-sided t-test (*** *P* < 0.001). Black bars indicate comparison. Mean, *P*-value, cell number indicated on plot. **D)** Scatter plot depicting significantly altered genes in *NFKB2*-KD cells. Significance score calculated by -Log2(*P*-value), only genes with significance score > 8 included in graph. (Red = increased expression, blue = decreased expression). Differential expression scoring calculated via *pyspade*. **E)** RT-qPCR data showing expression of *GAPDH*-normalized *NFKB2* expression and *GAPDH*-normalized *CXCL14* expression in scramble control or *NFKB2*-targeted shRNA-knockdown cells from undifferentiated control cultures (n=3). *P*-value calculated by Students t-test (** *P* < 0.01).

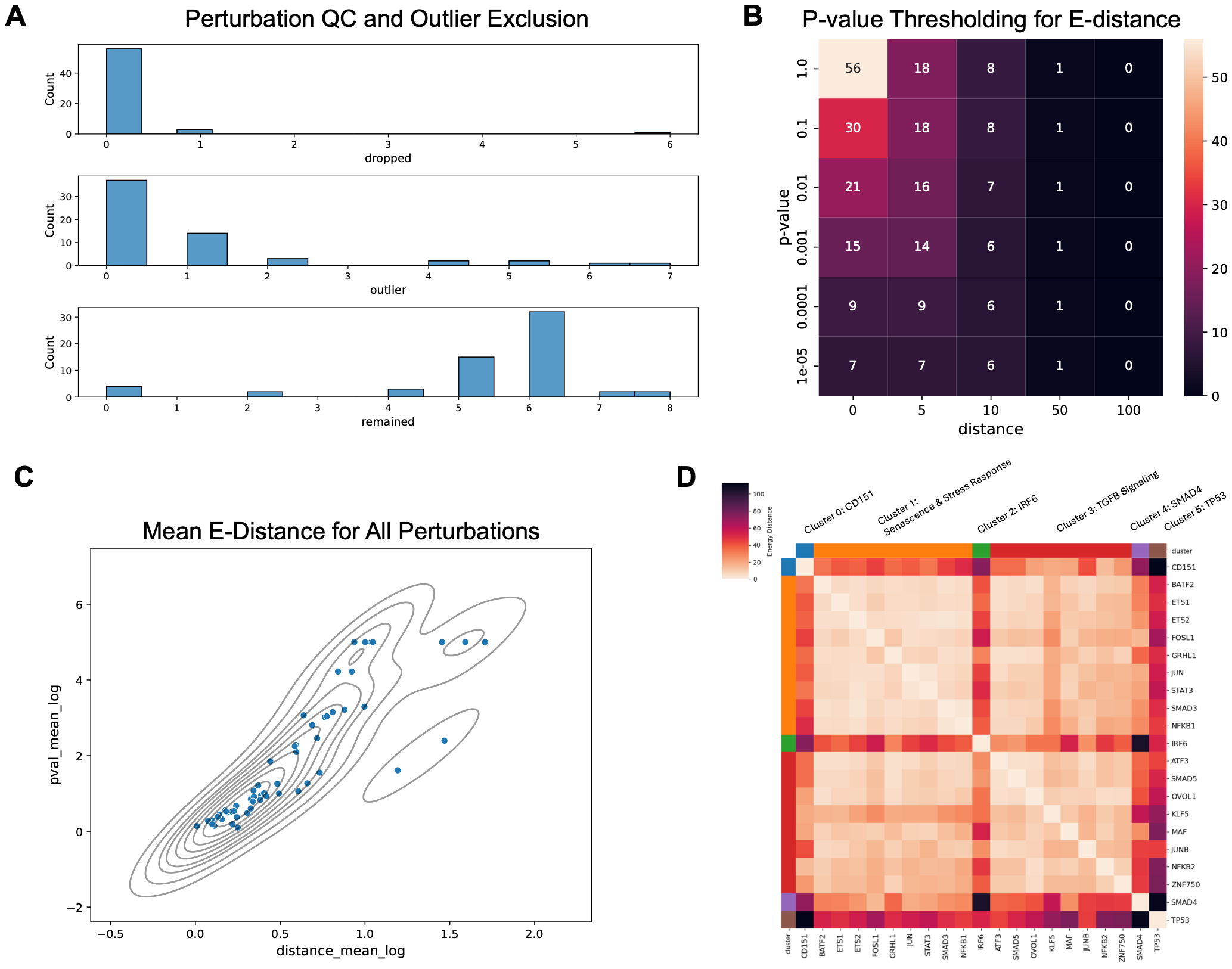

**Supplementary Figure 5) Energy distance QC and metrics. A)** Bar plots indicating the number of perturbations (sgRNA species) excluded from energy distance analysis. **B)** Heat map indicating number of perturbations (target-genes) at *P*-value and energy distance thresholds. Color corresponds to perturbation number as indicated on heat map. *P*-value of 0.001 was selected for further analysis. **C)** Scatter density plot depicting sgRNA *P*-values as a function of transcriptome distance (Log10 Energy Distance). Each point represents a single perturbation (target-gene). **D)** Heat-map depicting pairwise Energy Distance between perturbations (target-gene). Color represents pairwise distance as indicated on heat map. Clustered genes are grouped and labeled based upon top go enrichment terms, or by perturbation name if only including a single perturbation.

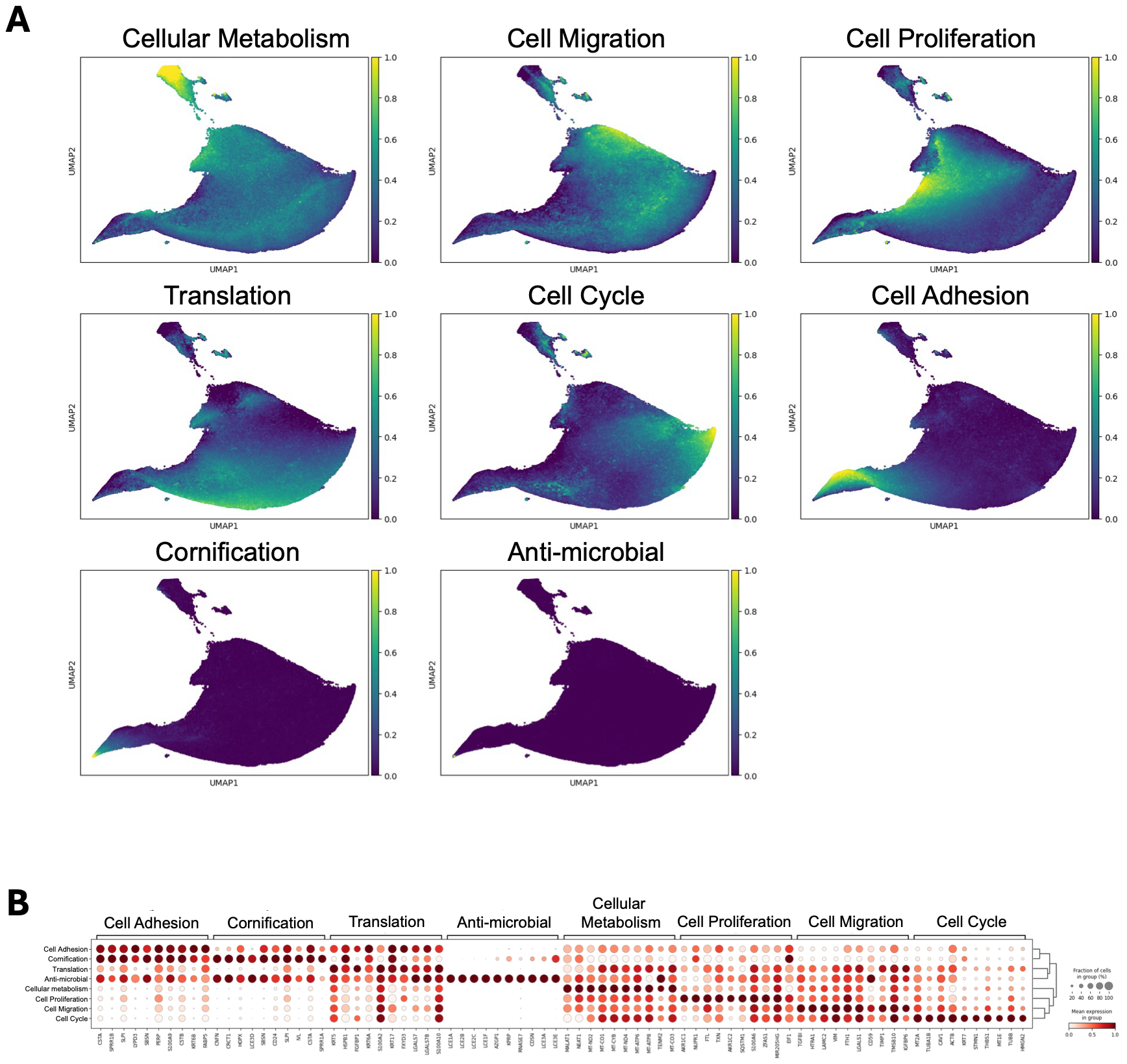

**Supplementary Figure 6) cNMF expression and ranked gene expression. A)** UMAP plots depicting cNMF calculated gene expression program (GEP) scores per cell. Each point represents one cell. Color represents expression score as indicated on plot. Each plot represents the score of one GEP in the entire population**. B)** Dot plot depicting top 10 ranked genes expressed in high GEP scoring cells. Cells segregated based on top GEP score per cell. Rank genes calculated using *scanpy* rank genes. GEP group and gene expression indicated on plot. Dot size corresponds to population percentage expressing the indicated gene. Color indicates normalized gene expression in the population.

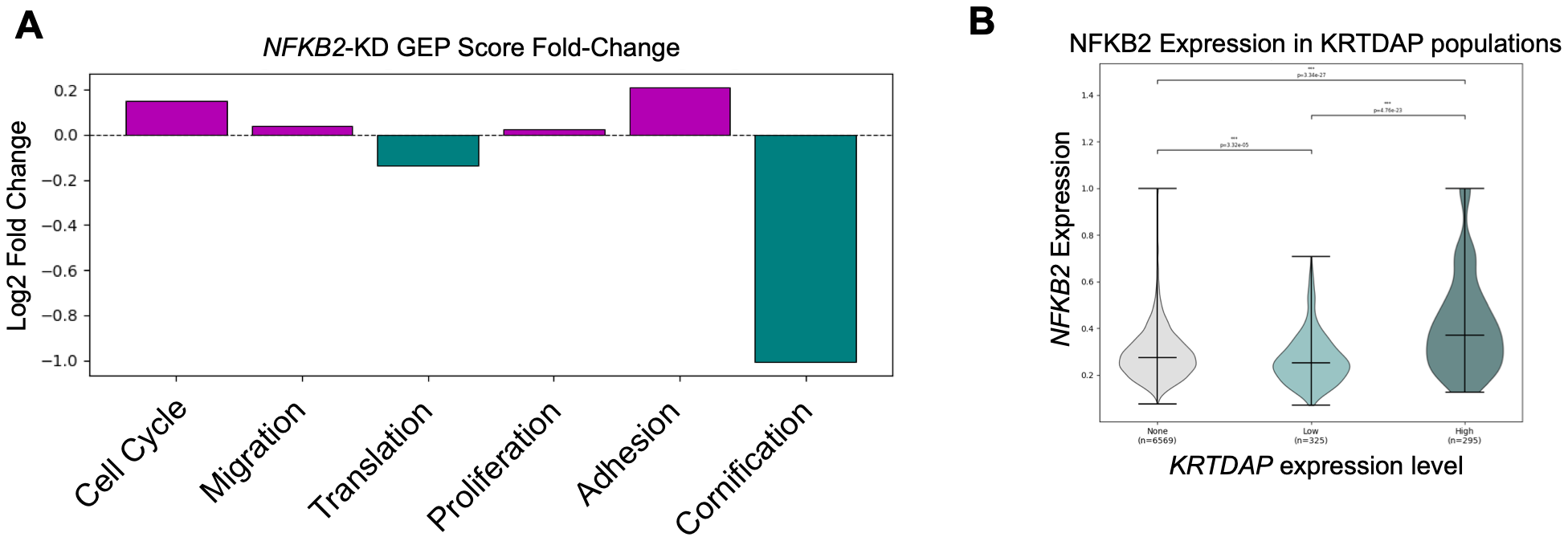

**Supplementary Figure 7) Transcription factor knockdown GEP scoring. A)** Bar plot depicting Log2 fold change of GEP scores in *NFKB2*-KD cells compared to non-targeting control cells. GEP identity indicated on graph, ordered by median pseudotime. **B)** Violin plot showing biphasic *GAPDH*-normalized *NFKB2* expression in basal-to-granular keratinocyte lineage of epidermal organoid. Error bars indicate mean ± SD. Cells segmented based upon *KRTDAP* expression level. Cells segregated at *KRTDAP* expression median. *P*-value calculated by Mann-Whitney two-sided t-test (*** *P* < 0.001). Black bars indicate comparison. Mean, *P*-value, cell number indicated on plot.

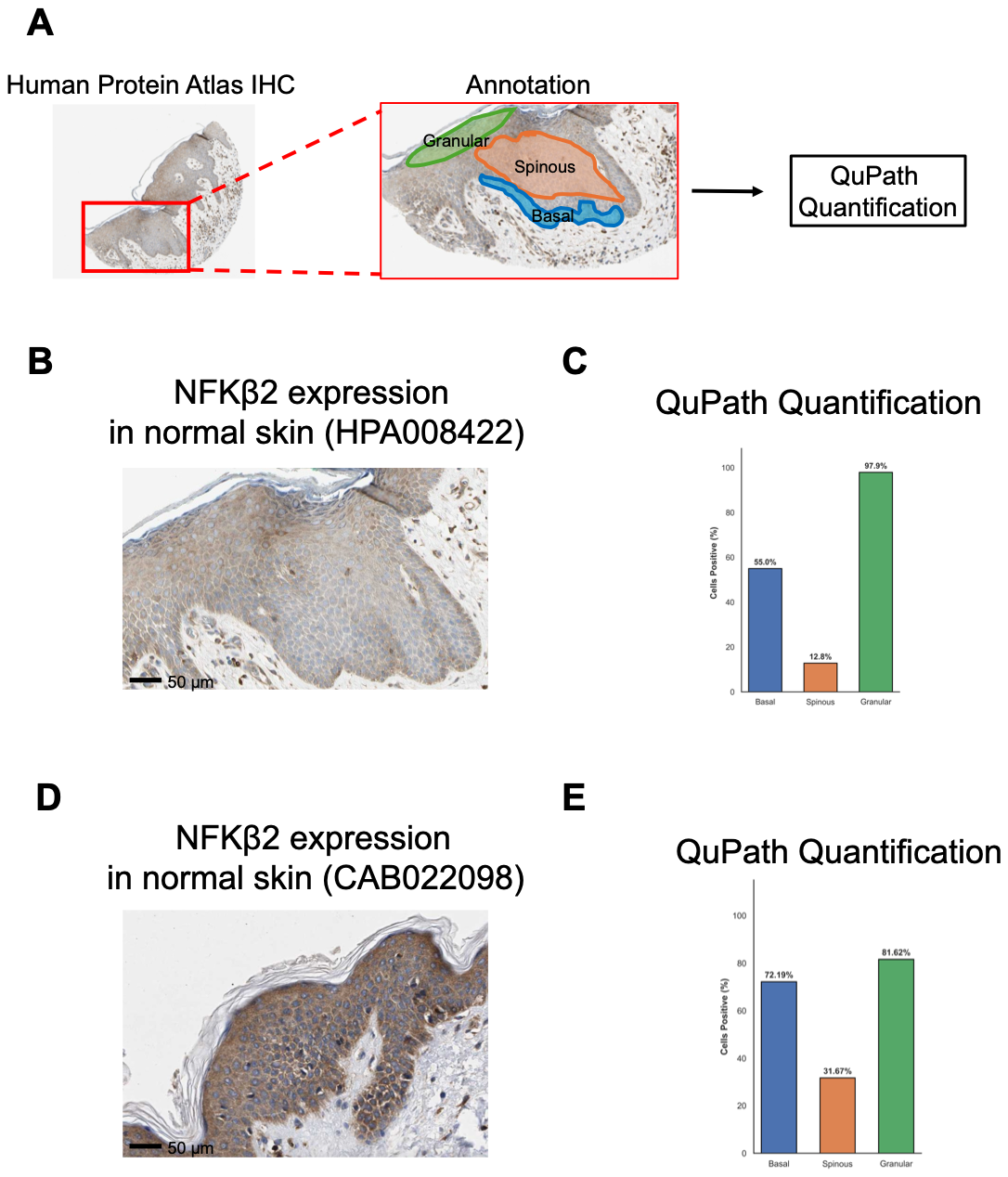

**Supplementary Figure 8) NFKB2 expression in normal human skin. A)** Schematic depicting annotation and quantification workflow for NFKB2 IHC analysis. **B)** Region of interest (ROI) for human protein atlas (HPA) sample HPA008422. **C)** QuPath cell percentage expressing NFKB2 in ROI for HPA sample CAB022098. **D)** ROI for HPA sample HPA008422. **E)** QuPath cell percentage expressing NFKB2 in ROI for HPA sample CAB022098.
